# Mast Cells Enhance Sterile Inflammation in Chronic Nonbacterial Osteomyelitis

**DOI:** 10.1101/259275

**Authors:** Stephanie Young, Namit Sharma, Jae Hoon Peter Lee, Violeta Chitu, Volker Neumeister, Elisabeth Sohr, E. Richard Stanley, Christian M. Hedrich, Andrew W.B. Craig

**Affiliations:** Department of Biomedical and Molecular Sciences, Queen’s University, Kingston, ON K7L 3N6 Canada; Department of Developmental and Molecular Biology, Albert Einstein College of Medicine, Bronx, NY 10461, USA; Departments of Clinical Chemistry and Laboratory Medicine, University Hospital Carl Gustav Carus, Technical University Dresden, Dresden, Germany; Pediatric Rheumatology and Immunology, Children’s Hospital Dresden, Technical University Dresden, Dresden, Germany 74, D-01307; Department of Women’s & Children’s Health, Institute of Translational Medicine, University of Liverpool, and Department of Paediatric Rheumatology, Alder Hey Children’s NHS Foundation Trust Hospital, Liverpool, UK

**Keywords:** autoinflammatory disorders, chronic nonbacterial osteomyelitis, chronic multifocal osteomyelitis, innate immune system, mast cells, cytokines and inflammation, osteoclasts

## Abstract

Chronic nonbacterial osteomyelitis (CNO) is an autoinflammatory bone disease. While some patients exhibit bone lesions at single sites, most patients develop chronically active or recurrent bone inflammation at multiple sites, and are then diagnosed with recurrent multifocal osteomyelitis (CRMO). Chronic multifocal osteomyelitis (CMO) mice develop IL-1β-driven sterile bone lesions reminscent of severe CRMO. Mechanistically, CMO disease arises due to loss of PSTPIP2, a negative regulator of macrophages, osteoclasts and neutrophils. The goal of this study was to evaluate the potential involvement of mast cells in CMO/CRMO disease pathophysiology. Here, we show that mast cells accumulate in the inflamed tissues from CMO mice, and mast cell protease Mcpt1 was detected in the peripheral blood. The role of mast cells in CMO disease was investigated using a transgenic model of connective tissue mast cell depletion (Mcpt5-Cre:Rosa26-Stop^fl/fl^-DTa) that was crossed with CMO mice. The resulting CMO/MC-mice showed a significant delay in disease onset compared to age-matched CMO mice. At 5-6 months of age, CMO/MC- mice had fewer bone lesions and immune infiltration in the popliteal lymph nodes that drain the affected tail and paw tissues. To test the relevance of mast cells to human CRMO, we tested serum samples from a cohort of healthy controls or CRMO patients at diagnosis. Interestingly, mast cell chymase was elevated in CRMO patients as well as patients with oligoclonal juvenile arthritis. Tryptase-positive mast cells were also detected in bone lesions from CRMO patients as well as patients with bacterial osteomyelitis. Taken together, our results identify mast cells as cellular contributors to bone inflammation in CMO/CRMO. Observations of this study promise potential for mast cells and derived mediators as future biomarkers and/or therapeutic targets.

## INTRODUCTION

Chronic nonbacterial osteomyelitis (CNO) is an autoinflammatory bone disease (1). While some patients exhibit bone lesions at single sites, most patients develop chronically active or recurrent bone inflammation at multiple sites, and are then diagnosed with recurrent multifocal osteomyelitis (CRMO) (2–6). CRMO partially resembles other syndromic autoinflammatory bone diseases, such as deficiency of the interleukin (IL)-1 receptor antagonist (DIRA) or Majeed syndrome (7, 8). CRMO patients with active disease are characterized by elevated pro-inflammatory cytokines (Tumor necrosis factor (TNF)-α, IL-1β, IL-6 and IL-8) in the serum (9), and reduced IL-10 production in peripheral blood monocytes (10). In CRMO patient samples, reduced IL-10 expression correlates with increased inflammasome expression and activation (11), leading to increased IL-1β-driven inflammatory bone loss (1, 12, 13).

Several key features of severe CRMO are modeled in chronic multifocal osteomyelitis (CMO) mice (3, 14). CMO mice carry destabilizing homozygous L98P mutations in proline-serine-threonine phosphatase interacting protein 2 (*Pstpip2*) (3, 14). Expression of PSTPIP2 is restricted to hematopoietic progenitors, and innate immune cells (14–17). PSTPIP2 suppresses differentiation and effector function of macrophages (14, 18, 19) neutrophils (16, 17, 20, 21) and osteoclasts (22). This may be explained by PSTPIP2 promoting plasma membrane recruitment of negative regulators of immune cell activation (23, 24), including PEST protein-tyrosine phosphatases (22, 25, 26), C-terminal Src kinase (Csk) (26), and the phosphatidylinositol-3, 4, 5-trisphosphate 5-phosphatase SHIP-1 (26). These interactions are enhanced in response to immune cell activation by a variety of receptors (18, 26, 27). PSTPIP2-deficient neutrophils show elevated responses to inflammatory stimuli (lipopolysaccharide (LPS), FcR capping, crystalline silica (CS)) (16, 26), likely due to increased Src/Ras/Erk (28, 29), and PI3K/Akt signaling (30, 31). However, PSTPIP2 is also expressed in other cells, including mast cells (14), but no functional studies have been reported to date.

Mast cells are specialized innate immune cells residing near potential portals of entry for pathogens (32). Mast cell degranulation results in the release of mast cell proteases (chymase, tryptase), cytokines/chemokines (TNF-α, CXCL1/KC), and histamine that all promote tissue inflammation, leukocyte recruitment, and blood vessel permeability (33, 34). Mast cell activation also leads to rapid production and release of lipid mediators implicated in leukocyte recruitment and tissue inflammation (35, 36). Studies using transgenic models of connective tissue mast cell (CTMC) ablation implicated key roles of mast cells in allergic skin inflammation and arthritis (37–39). Hence, mast cells and their mediators may contribute to inflammatory disorders driven by aberrantly activated innate immune cells, such as CMO/CRMO.

In this study, we demonstrate that mast cells accumulate in inflammatory bone lesions in CMO mice, and that genetic ablation of CTMCs results in delayed onset and reduced severity of the inflammatory disease. Furthermore, we detect mast cell infiltrates in bone tissue lesions of CRMO patients, and the release of mast cell chymase in the serum of CRMO patients at diagnosis.

## MATERIALS AND METHODS

### Mice

The constitutive CTMC ablation model (Mcpt5-Cre:Rosa-Stop^fl/fl^-Dta) (37) was crossed for 10 generations with *Pstpip2*^cmo/cmo^ (CMO) mice (22) to achieve a uniform BALB/cAnPt background for all (CMO/MC-) mice. All animals were housed at Queen’s University Animal Care Services under specific pathogen-free conditions. All procedures were approved by the Queen’s University Animal Care committee in accordance with Canadian Council on Animal Care guidelines.

### Disease scoring, pharmacological treatments, and preparation of tissue homogenates

Wild-type (WT), CMO and CMO/MC- mice were monitored for the initial appearance of a tail kink or paw deformity to recored the date of disease onset. For tissue cytokine measurements, equal weights of distal tail segments and hind paws were snap frozen, homogenized, and ELISA performed for IL-1β (e-bioscience). To distinguish levels of pro-IL-1β and mature IL-1β, some homogenates were analyzed by immunoblot using mouse anti-IL-1β (3A6, Cell Signaling Technology; anti-β-actin (Santa Cruz Biotechnology) served as a loading control).

### Lymph node immune profiling

Popliteal lymph nodes from each mouse were collected, weighed, and manually homogenized in PBS-EDTA solution. Cells were counted, resuspended in PBS with sodium azide (0.1%) and BSA (0.5%), and stained with: FITC anti- CD117 (c-KIT, Biolegend), PE anti- FcεRI (Biolegend), FITC anti-I-Ek/RT1-D (MHC II, BioLegend), FITC anti-Ly6C (BD Biosciences), and PE anti-Ly6G (BD Biosciences). Samples were analyzed using a cytomics Fc500 (Beckman Coulter), and FlowJo software (Tree Star), with gating set to exclude dead cells (low forward and side scatter; isotype control antibodies were used to set threshold values).

### Micro-computed tomography

Tail and hind paws harvested from male mice at 5 months of age were analyzed in a Skyscan 1172 high-resolution micro-CT scanner (Bruker, Milton, Canada). With the X-ray source at 100 kVp and 100 μA, and the use of a 0.5 mm aluminium filter, each specimen was rotated 360° around the vertical axis to generate 1200 views in 5 h. The image projections were reconstructed into digital cross-sections using the Feldkamp algorithm (40) for cone beam CT. The resulting 3D data block contained 2000×1000×1000 voxels of 13.4 μm, and resampled to match the resolution (28 μm) and data size of the 3D segmented atlas. The down-sampling of the 3D images was performed as the 2× decrease in voxel size allowed for an 8× reduction in the amount of RAM and image processing times. The volume fraction data (BV/TV) for serial sections was analyzed with Image J software plugin BoneJ (41).

### CMO mouse tissue staining

Tissues collected from diseased mice were placed in neutral buffered formalin. Tail and hind paw tissues were decalcified in 20% EDTA (biweekly refreshed for >2 months) prior to paraffin embedding. Tissue sections (4 μm thickness) were stained with either H&E, Alcian Blue staining of mast cells (42), or tartrate-resistant acid phosphatase (TRAP) staining of osteoclasts (22).

### Human CRMO patient serum and bone biopsy analyses

The cohort of CRMO patient serum samples taken at initial diagnosis was described previously (9). Chymase levels were measured by ELISA (CUSABIO) at University Hospital Carl Custav Carus in sera from 20 CRMO patients and 21 matched controls. Patients and controls had no records of allergies in their patient records, and all patients and/or their legal guardians gave written informed consent (ethics committees at University of Würzburg and University of Technology Dresden) (9). Formalin-fixed, decalcified, and paraffin-embedded bone biopsy specimen from patients with bacterial osteomyelititis (OM; n=3), early CRMO lesions (n=4), chronic phase CRMO lesions (n=4), or healthy control osteotomies (n=2) were stained with a human tryptase antibody (Abcam, ab2378, dilution: 1:500) using standard techniques. Nuclei were stained using Hämatoxylin, and Bluing Reagent on the BenchMarkXT staining system (Ventana Medical Systems/Roche). Slides were scanned on the AxioScan Z1 (Zeiss, Germany), and quantified using Zen Blue software (Zeiss, Germany) in 15-25 fields of view (32188.774 μm^2^).

### Statistical analysis

When comparing three genotypes, nonparametric Kruskal-Wallis one-way ANOVA, followed by a Dunn’s multiple comparison test was used. When comparing two genotypes, the Student’s t test was used (GraphPad Software, La Jolla, Calif). All graphs represent mean ± standard deviation.

## RESULTS

### Evidence of CTMC activation and role in chronic multifocal osteomyelitis

To test the relevance of CTMCs to CMO disease pathophysiology, we collected tail tissues from 5 month old wild-type (WT) and *Pstpip2*^*cmo/cmo*^ (CMO) mice, when severe multifocal osteomyelitis develops (3, 16, 43). Compared to baseline levels of Alcian blue-stained CTMCs in WT tail tissue sections, we observed a trend towards increased CTMCs in CMO tail tissues (**Figure 1A, B**). To test for mast cell activation and mediator release, we collected mouse plasma at 5 months of age, and measured the levels of mouse mast cell protease-1 (MCPT1) by ELISA. MCPT1 levels were significantly elevated in plasma from CMO mice, as compared to WT animals (**Figure 1C**). These results suggest that CTMCs are present within inflamed tissues of CMO mice and that mast cell mediators are produced in this autoinflammatory disease model.

**FIGURE 1.**
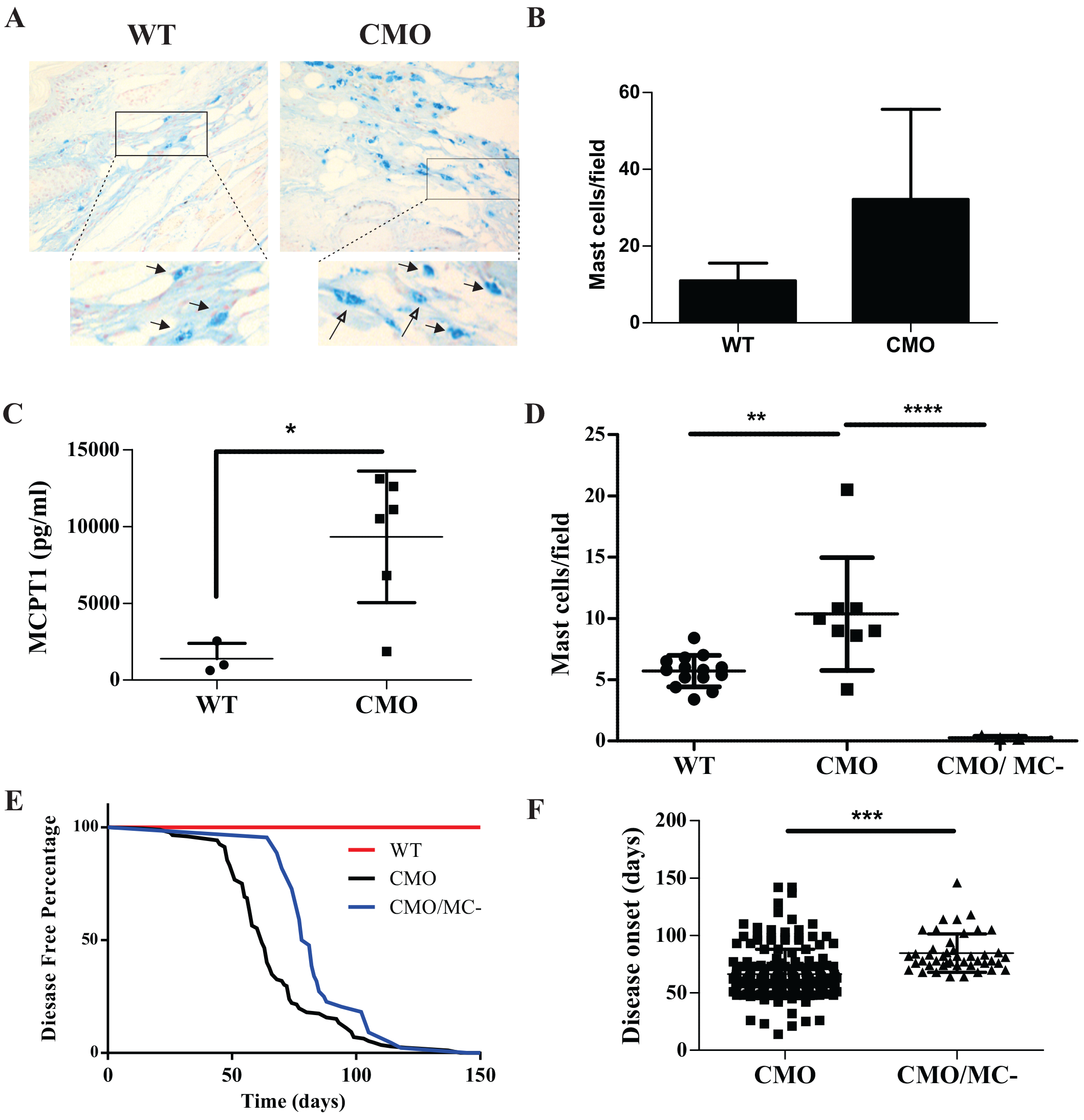
CTMCs promote CMO disease onset. (**A)** Representative images of mast cells detected with Alcian blue staining of decalcified tail tissue sections from 5 month old WT and CMO male mice (insets show magnified view with arrows indicating mast cells). (**B)** Quantification of mast cell density in WT and CMO tail tissue sections stained with Alcian blue. Graph depicts the average mast cell density per field (mean ± SD, 5 fields/sample, 3 mice/group). (**C)** Levels of MCPT1 were profiled in plasma from 5 month old male WT (N=3) and CMO (N=6) mice by ELISA. (**D)** Graph depicts quantification of mast cell populations in Alcian blue-stained ear tissue sections (N=6 per group; 5 fields/mouse; * P<0.05). (**E**) Kaplan-Meier disease-free survival curve is shown for WT (N=50), CMO (N=172), and CMO/MC- (N=42) mice. (**F)** Graph depicts the average age of disease onset in CMO (N=40) and CMO/MC- (N=43) mice (*** P<0.001).

To test the contribution of CTMCs to the development of CMO disease, we employed a transgenic model that results in constitutive ablation of CTMCs due to Diptheria toxin-α expression in mature CTMCs (Mcpt5-Cre:Rosa26-Stop^fl/fl^-DTa) (37). Both transgenes were crossed with CMO mice (3), and backrossed >9 generations to produce CTMC-deficient CMO animals (hereafter referred to as CMO/MC-) on a uniform Balb/c background. We compared CTMC densities in ear skin tissue sections from WT, CMO and CMO/MC- mice by Alcian blue staining. Compared to WT, we observed elevated CTMC density in CMO ear skin lesions (**Figure 1D**). CMO/MC- ear skin was largely devoid of CTMCs (**Figure 1D**), and this also correlated with reduced inflammation and edema (**Figure S1**). These results suggest a potential role for CTMC activation in CMO disease, and demonstrates the efficacy of mast cell ablation in the CMO/MC-model.

To determine the role of CTMCs in CMO disease onset, cohorts of age-matched CMO and CMO/MC- mice were monitored for early symptoms of CMO, including tail kinks or deformities in hind paws. The CMO/MC- cohort showed a significant delay in disease onset compared to CMO mice, while WT mice showed no disease (**Figure 1E, F**). Overall, these results implicate mast cells in promoting inflammation in the CMO model.

### CTMC deficiency protects against bone damage in CMO mice

To test the role of CTMCs in CMO disease progression, tail and paw tissues were collected from cohorts of male WT, CMO and CMO/MC- mice at 5 months of age. Analysis of these tissues and microcomputed tomography (μCT) scans revealed fewer deformities and bone lesions within the tails and paws of CMO/MC- mice compared to CMO mice (**Figure 2A**). Quantification of bone density from the μCT scans of 6 mice from each group revealed that the CMO/MC- cohort were significantly protected from bone loss observed in CMO mice (**Figure 2B**). This also correlated with fewer TRAP-positive osteoclasts in CMO/MC- compared to CMO tail tissue sections (**Figure S2**). These results indicate that CTMCs promote bone inflammation and the resulting damage by osteoclasts that are hallmarks of this autoinflammatory disease.

**FIGURE 2.**
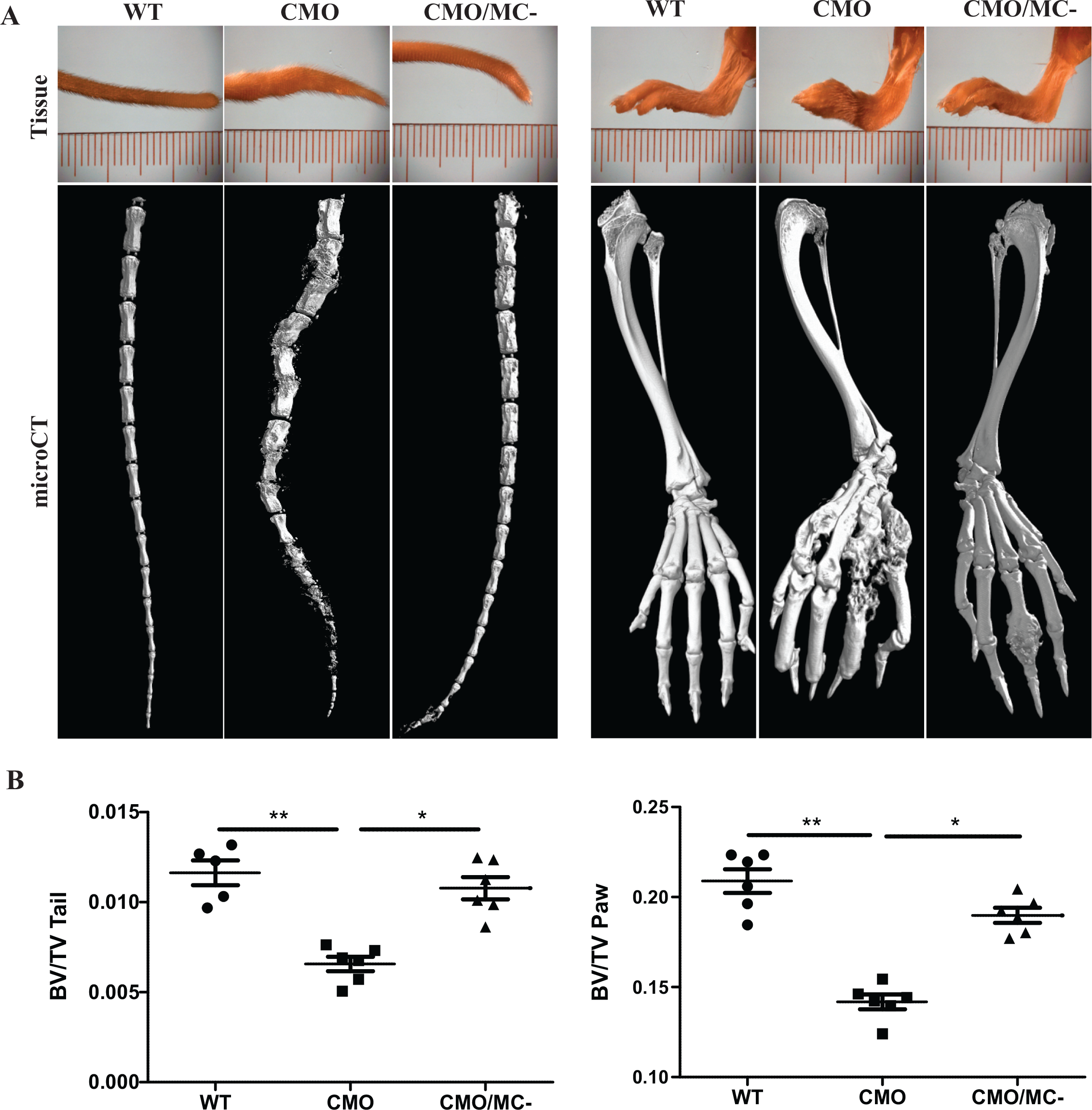
CTMCs promote bone lesion severity in CMO mice. (**A)** Representative images of tails and hind paw tissues are shown, along with μCT images of underlying bone lesions in male 5 month old WT, CMO and CMO/MC- mice. (**B)** Graphs depict quantification of bone volume to total volume ratios for each stack of μCT images of tails and hind paws for each cohort (N=6 per group; *P<0.05, **P<0.01).

### CTMCs promote IL-1β production and immune cell infiltration in CMO mice

Previous studies of CMO mice revealed a critical role for elevated IL-1β production and signaling in this model (16, 43). To test how CTMCs may contribute to the IL-1β production, we prepared tail tissue homogenates from 5 month old WT, CMO and CMO/MC- mice, and measured IL-1β levels by ELISA. Consistent with previous studies (16, 43), CMO tissues had significantly higher IL-1β levels compared to WT mice, but these levels were largely normalized in CMO/MC- mice (**Figure 3A**). Using immunoblot assays to separate the pro-IL-1β and mature IL-1β, we show that CMO/MC- tail tissues had less of the mature form of IL-1β (**Figure 3B**). These results suggest that CTMCs either contribute to IL-1β production in the CMO model, or the recruitment of other immune cells that produce IL-1β in this model.

**FIGURE 3.**
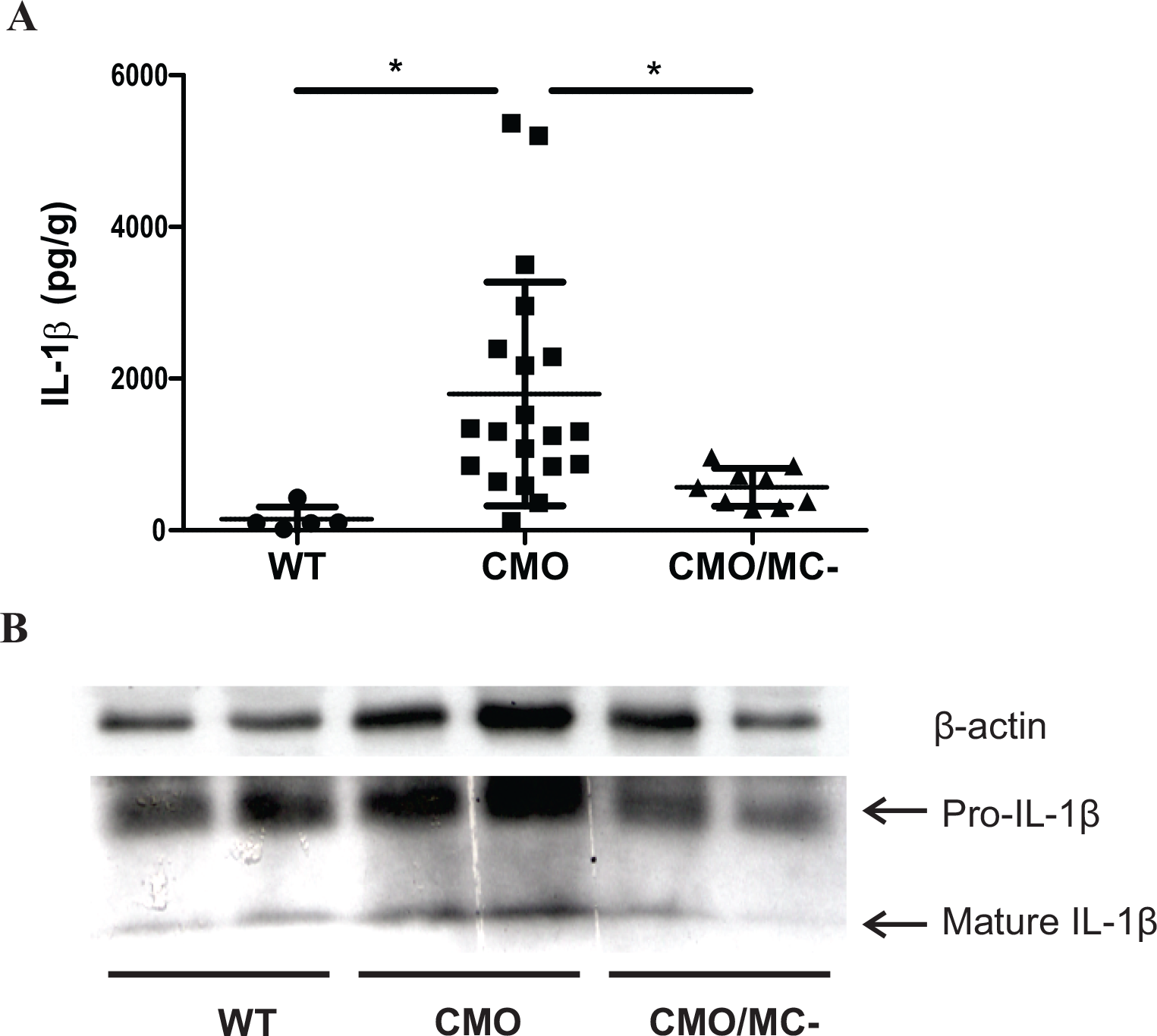
CTMC deficiency impairs IL-1β production in CMO disease tissue. **(A)** Graph depicts the levels of IL-1β detected by ELISA using tail tissue homogenates from 5 month old male WT (N=5), CMO (N=18), and CMO/MC- mice (N=9). (**B**) Respresentative immunoblot detection of pro-IL-1β and mature IL-1β in homogenates of tail tissues from WT, CMO and CMO/MC- male mice at 5 months of age. The positions of pro-IL-1β and mature IL-1β are shown, with β-actin used as a loading control. (*P<0.05)

Prior studies correlated the severity of CMO with the size of popliteal lymph nodes that “drain” the tail and hind paws (14, 17). Indeed, popliteal lymph node mass was elevated in CMO mice as compared to WT animals, but this expansion was reduced in CMO/MC- mice (**Figure 4A**). Flow cytometric analyses of immune cell markers in lymph nodes revealed that CMO mice had significantly elevated numbers of FcεRI^+^/KIT^+^ mast cells, MHCII^+^ activated immune cells, and a trend towards increased Ly6G^+^/Ly6C^+^ (GR-1) myeloid cells in lymph nodes compared to either WT or CMO/MC- mice (**Figure 4B**). Together, these results support increased mast cell density in CMO disease tissues, and a role of CTMCs in the recruitment of immune cells in CMO mice.

**FIGURE 4.**
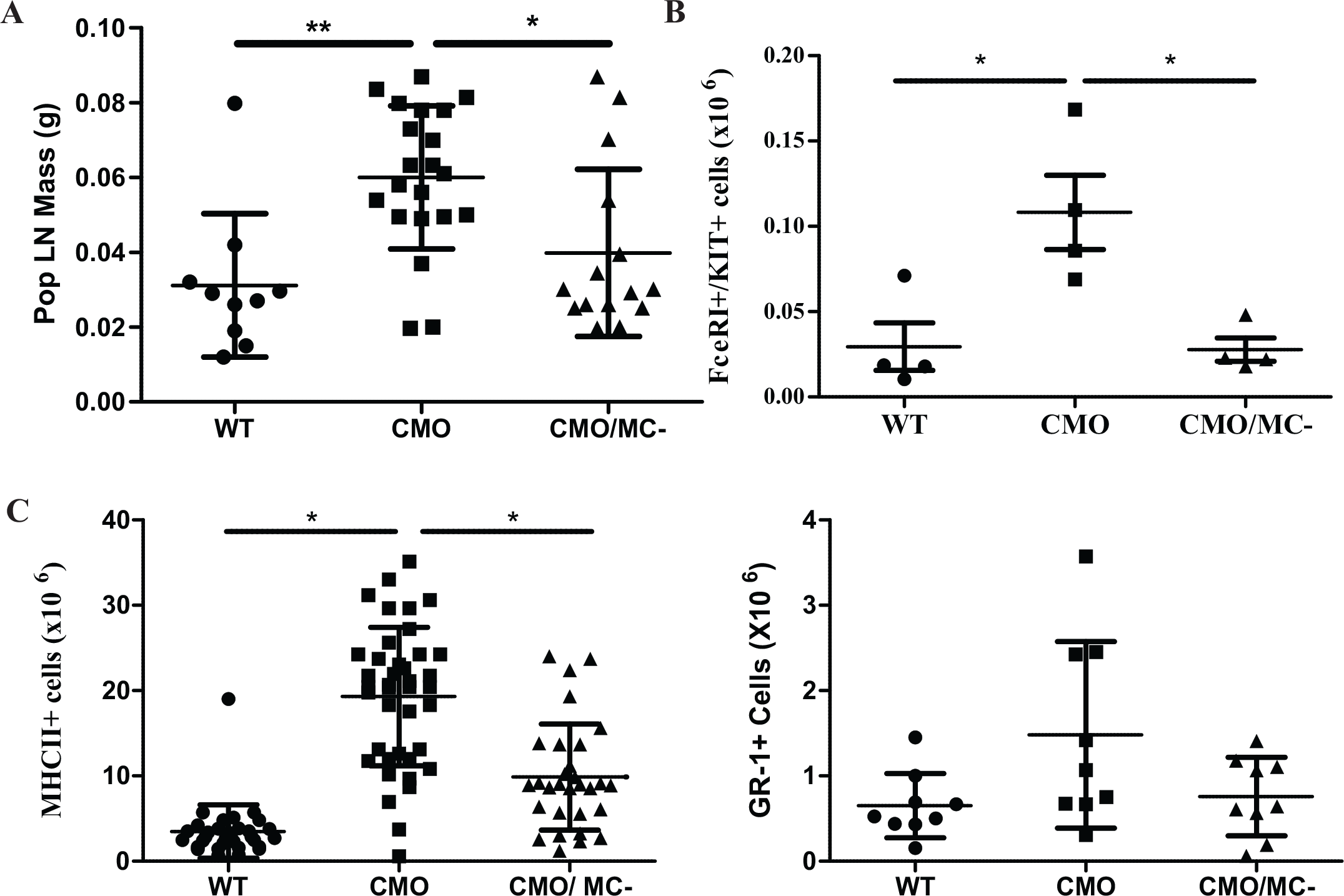
Impaired immune cell recruitment to popliteal lymph nodes in CMO/MC- mice. **(A)** Graph depicts the mean mass of popliteal lymph nodes for WT, CMO, and CMO/MC- male mice (N=10-20/group; (*P<0.05, **P<0.01). (**B)** Dot plot depicts the absolute numbers of mast cells (FcεRI^+^/KIT^+^) in popliteal lymph nodes from WT, CMO and CMO/MC- male mice based on flow cytometry (N=4/group). (**C)** Dot plots depict the absolute numbers of MHCII^+^ and GR-1+ immune cells in popliteal lymph nodes from WT, CMO and CMO/MC- male mice based on flow cytometry (N=9-21/group; results pooled from repeated experiments, *P<0.05).

### Mast cell activation in CRMO patient samples

To test the potential involvement of mast cells in human CRMO, we screened sera from a previously reported cohort of treatment naïve, newly diagnosed CRMO patients, oligoarticular juvenile arthritis (Oligo JIA) patients, and healthy controls (9). We tested for mast cell chymase by ELISA, and detected very low levels of chymase in 4 of 21 healthy controls, whereas the vast majority of CRMO patients (17 of 20) exhibited detectable serum chymase levels (**Figure 5A**). No patients in these cohorts had reported allergies. Of note, a comparable increase in serum chymase levels was also observed in Oligo JIA patients (**Figure 5A**), which is consistent with a recent study implicating mast cells in arthritis disease models (39).

**FIGURE 5.**
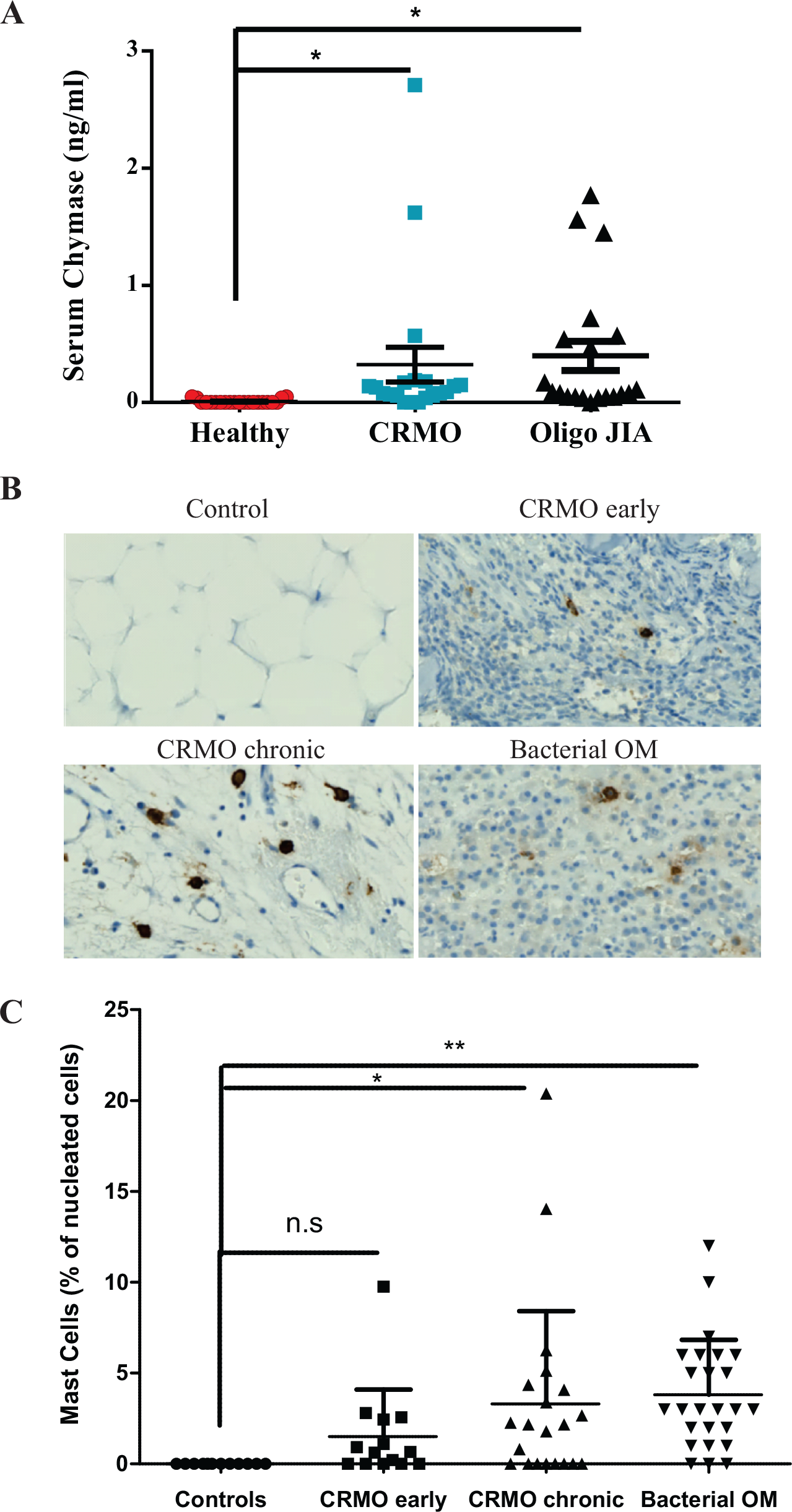
Detection of mast cells and mast cell mediators in CRMO patient samples. (**A)** Serum samples from human patients with CRMO (N=20), oligoarticular JIA patients (N=20), or from healthy controls (N=21) were tested for the levels of mast cell chymase by ELISA. Dot plot depicts individual values with median and interquartile range overlayed. (**B)** Representative images of tryptase staining of bone from healthy controlscontrol, early and chronic CRMO patients and infectious osteomyelitis patients. (**C)** Dot plot depicts the percentage of tryptase-positive mast cells relative to total nucleated cells in the field of view for bone sections from healthy controls, early and chronic lesions from CRMO patients, and bacterial OM patients.

To assess mast cell infiltration to inflamed bone tissue, we performed immunohistochemistry staining of tryptase-positive mast cells in tissue sections from bone biopsies taken from healthy controls (osteotomies), CRMO patients, or bacterial osteomyelitis (OM) patients. While no mast cells were detected in bone biopsies from healthy individuals, we detected mast cells in CRMO lesions, including early CRMO lesions marked by innate immune infiltrates (**Figure 5B,C**). Particularly high mast cell counts were detected in chronic CRMO lesions marked by coexisting infiltrates of innate immune cells and lymphocytes (**Figure 5B,C**). Mast cell counts were also increased in bacterial OM bone biopsies when compared to controls (**Figure 5B,C**).

## DISCUSSION

Studies in CMO mice and related mouse models have provided insights into the pathophysiology of human CRMO, a rare autoinflammatory disease. This includes the identification of a skewed microbiome, increased IL-1β production, and aberrantly activated innate immune cells (14, 16, 21, 22, 43). In the study presented here, we show that mast cells accumulate in CMO lesions, and promote the accumulation of bone inflammation and lesions. By crossing CMO mice with CTMC-deficient animals (37), we provide evidence for CTMCs promoting CMO disease onset and severity. We also translate these studies to human CRMO, by providing evidence of mast cell infiltrates in bone biopsies from CRMO patients, and elevated levels of mast cell chymase in the serum of CRMO patients at diagnosis. Together, these findings implicate mast cells in promoting bone inflammation in CMO mice, and suggest a role for mast cells in the pathophysiology of CRMO in humans. Our model depicts several candidate mediators from mast cells that impinge on recruitment and activation of the innate immune cells and osteoclasts that trigger autoinflammation (**Figure 6**).

**FIGURE 6.**
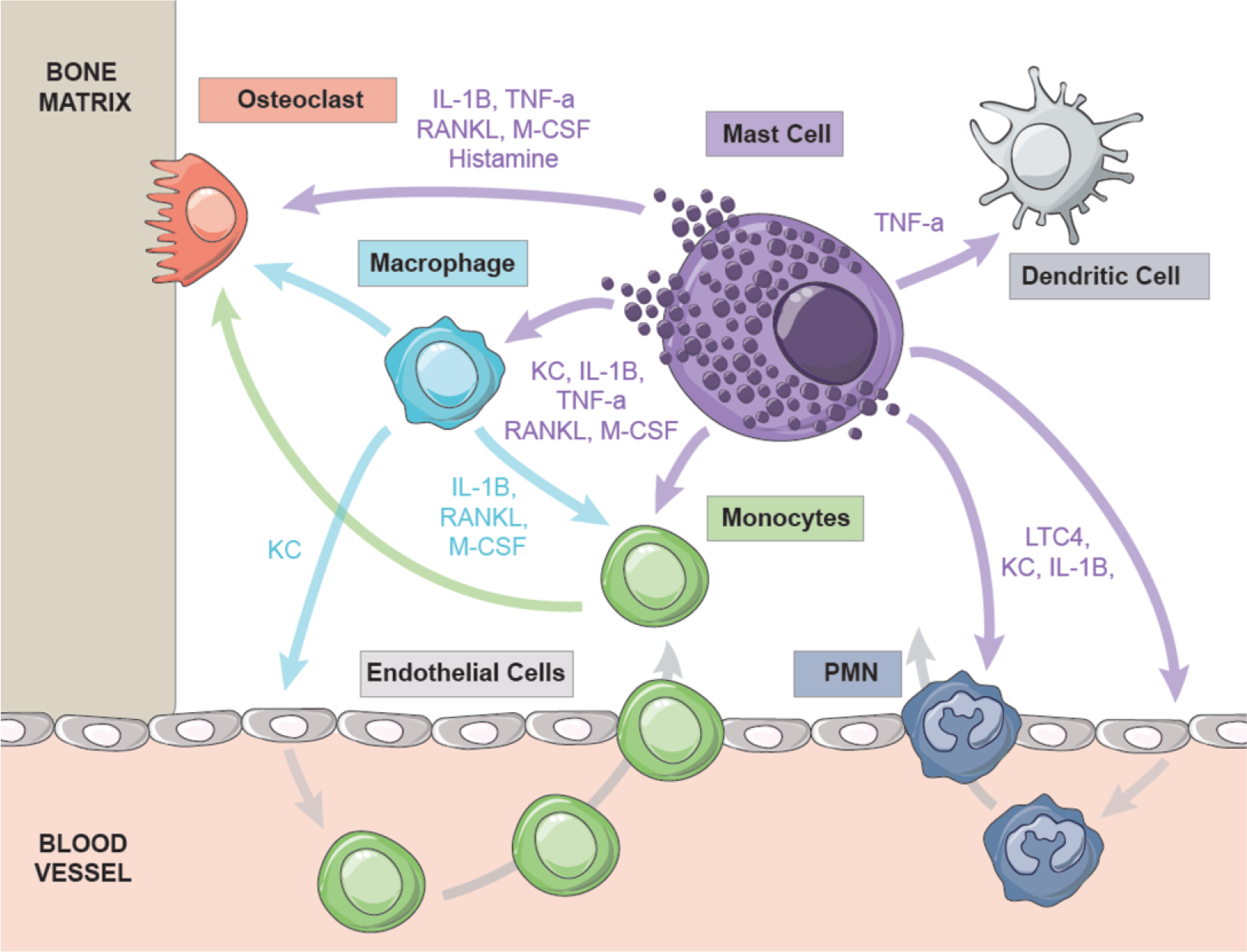
Hypothetical model of potential crosstalk between mast cell mediators and cell types implicated in autoinflammation. Model depicts the effects of mast cell mediators on recruitment and activation of innate immune cells that are previously implicated in promoting IL-1β-driven sterile inflammation (see text for details).

A recent study provides evidence of functional cross-talk between mast cells and osteoclasts, with mast cell degranulation promoting osteoclast differentiation and activation (44). Our study provides further *in vivo* evidence of mast cell activation contributing to increased density of osteoclasts and the extent of bone loss in the CMO model. Future studies are warranted to fully dissect the potential cross-talk between mast cells, osteoclasts and other relevant cell types in CRMO and related diseases.

Since CMO mice are protected from disease when crossed onto either IL-1R-null or IL-1β-null backgrounds (16, 43), mast cells may be an important source of IL-1β. Alternatively, reduced IL-1β levels in CMO/MC- mice may also be due to impaired recruitment of IL-1β-producing immune cells. Indeed, draining lymph nodes from CMO/MC- mice exhibited significant reduction in the numbers of innate immune cells when compared to CMO mice. Mast cell proteases released in response to activation may also enhance cleavage of pro-IL-1β in inflamed tissues, based on studies in other models (45, 46). Interestingly, chymase is elevated in serum from CMO mice and CRMO patients at diagnosis. Thus, chymase may indeed participate in CMO and CRMO pathophysiology, and is worthy of further functional studies.

Previous studies suggested that neutrophils in CMO produce aberrant levels of IL-1β in an inflammasome-independent manner, which may be responsible for driving the CMO disease pathophysiology (16, 21). Indeed, the depletion of neutrophils using a Ly6G antibody protected CMO mice from bone lesion development (21). Here, we also detected significant elevation of Ly6C^+^/Ly6G^+^ cells in the CMO disease tissue sections compared to WT or CMO/MC- mice. This suggests that the Ly6C^+^/Ly6G^+^ cells may be recruited to the affected tissues prior to differentiation into tissue resident cells. It is worth noting that granulocytic myeloid-derived suppressor cells (G-MDSCs) express both Ly6C and Ly6G markers (Ly6C^lo^/Ly6G^hi^), and are recruited to the bone as the main osteoclast precursor population (47, 48). These cells then differentiate into osteoclasts in chronic inflammatory disorders such as arthritis and bone cancer (47–50). Thus, “neutrophil” depletion with inactivating anti-Ly6G antibodies in CMO mice (21) may have also depleted osteoclast precursor G-MDSCs, which would lead to fewer osteoclasts and reduced bone resorption. Regardless of how this data is interpreted, we observed a positive role of mast cells in recruitment of immune cells, and potential osteoclast precursors (G-MDSCs) as well (data not shown). Interestingly, MDSC proliferation has been reported to be mast cell dependent (51). Further studies using more selective tools to deplete and phenotype the inflammatory cells in this model are needed to better define CMO disease pathophysiology.

The identity of potential triggers of CMO disease, and restrictions of the phenotypes to certain tissues remains poorly defined. A previous study identified a skewed microbiome in CMO mice that leads to priming of neutrophils to express IL-1β (21). Based on the selective loss of CTMCs in our model, with no effect on mucosal mast cells (37, 42), we have not yet tested the mast cell involvement at the interface with a skewed microbiome. However, future studies using Cpa3^Cre^ mice, lacking both mucosal and connective tissue mast cells (52), would be an improved model to test the overall contribution of mast cells to this disease.

To translate findings from our study on the involvement of mast cells in murine CMO to CRMO in humans, we stained bone samples from CRMO patients and detected increased mast cell populations in CRMO patients. Profiling of CRMO serum biomarkers delivered a number of cytokines and chemokines that were elevated at diagnosis (9), with several of these known to be released by activated human mast cells (53–55). Our detection of elevated levels of mast cell chymase in sera from most CRMO patients provides new evidence of mast cell activation in this disease. This was also true for Oligo JIA patient sera, suggesting a common immune mechanism involving mast cells in these diseases. Of note, chymase can directly promote IL-1β production (56), and other mast cell mediators may also contribute to IL-1β-driven disease in CRMO patients (9, 12). This is also consistent with elevated mast cell density in bone tissue sections from CRMO patients compared to healthy controls that we report in this study. Thus, mast cells may amplify inflammation in the CMO model. This makes mast cells a promising source of new biomarkers and a potential target for new treatments.

In conclusion, we demonstrate a role for mast cells in the progression of autoinflammatory bone disease using mouse models and human CRMO patient samples. We show that aberrant mast cell activation enhances the cycle of IL-1β-driven sterile inflammation that triggers bone loss in this disease (**Figure 6**). Indeed, there is growing evidence of mast cell involvement in autoinflammation in a variety of models and diseases (57). This study provides rationale to study the effects of mast cell-targeted therapies in CRMO and related autoinflammatory bone diseases.

## AUTHOR CONTRIBUTIONS

SY, NS, JHPL, VN, and ES conducted experiments designed by CMH and AWBC. VC and ERS provided key resources and pilot experiments. The paper was written by AWBC, all authors contributed to editing, and AWBC had overall responsibility for the study.

## ACKNOWLEDGMENTS

We thank Matt Gordon at the Queen’s University Biomedical Imaging Centre (QUBIC) for help with flow cytometry, Queen’s Lab for Molecular Pathology (QLMP) for help with tissue embedding and histology, and Lisa Yu and Mark Henkelman at the Mouse Imaging Centre (mICE, Toronto) for performing microCT imaging.

## REFERENCES

1. Zhao Y, Ferguson PJ. Chronic Nonbacterial Osteomyelitis and Chronic Recurrent Multifocal Osteomyelitis in Children. Pediatric clinics of North America. 2018;65(4):783–800.

2. Ferguson PJ, Sandu M. Current understanding of the pathogenesis and management of chronic recurrent multifocal osteomyelitis. Current rheumatology reports. 2012;14(2):130–41.

3. Ferguson PJ, Bing X, Vasef MA, Ochoa LA, Mahgoub A, Waldschmidt TJ, et al. A missense mutation in pstpip2 is associated with the murine autoinflammatory disorder chronic multifocal osteomyelitis. Bone. 2006;38(1):41–7.

4. Ferguson PJ, Laxer RM. New discoveries in CRMO: IL-1beta, the neutrophil, and the microbiome implicated in disease pathogenesis in Pstpip2-deficient mice. Seminars in immunopathology. 2015;37(4):407–12.

5. Schnabel A, Range U, Hahn G, Siepmann T, Berner R, Hedrich CM. Unexpectedly high incidences of chronic non-bacterial as compared to bacterial osteomyelitis in children. Rheumatology international. 2016;36(12):1737–45.

6. Hofmann SR, Schnabel A, Rosen-Wolff A, Morbach H, Girschick HJ, Hedrich CM. Chronic Nonbacterial Osteomyelitis: Pathophysiological Concepts and Current Treatment Strategies. The Journal of rheumatology. 2016;43(11):1956–64.

7. Aksentijevich I, et al. An autoinflammatory disease with deficiency of the interleukin 1-receptor antagonist. The new England Journal of Medicine. 2009;360:2426–37.

8. Ferguson PJ, Chen S, Tayeh MK, Ochoa L, Leal SM, Pelet A, et al. Homozygous mutations in LPIN2 are responsible for the syndrome of chronic recurrent multifocal osteomyelitis and congenital dyserythropoietic anaemia (Majeed syndrome). Journal of medical genetics. 2005;42(7):551–7.

9. Hofmann SR, Kubasch AS, Range U, Laass MW, Morbach H, Girschick HJ, et al. Serum biomarkers for the diagnosis and monitoring of chronic recurrent multifocal osteomyelitis (CRMO). Rheumatology international. 2016;36(6):769–79.

10. Hofmann SR, Schwarz T, Moller JC, Morbach H, Schnabel A, Rosen-Wolff A, et al. Chronic non-bacterial osteomyelitis is associated with impaired Sp1 signaling, reduced IL10 promoter phosphorylation, and reduced myeloid IL-10 expression. Clinical immunology. 2011;141(3):317–27.

11. Brandt D, Sohr E, Pablik J, Schnabel A, Kapplusch F, Mabert K, et al. CD14(+) monocytes contribute to inflammation in chronic nonbacterial osteomyelitis (CNO) through increased NLRP3 inflammasome expression. Clinical immunology. 2018.

12. Hofmann SR, Kubasch AS, Ioannidis C, Rosen-Wolff A, Girschick HJ, Morbach H, et al. Altered expression of IL-10 family cytokines in monocytes from CRMO patients result in enhanced IL-1beta expression and release. Clinical immunology. 2015;161(2):300–7.

13. Cox AJ, Ferguson PJ. Update on the genetics of nonbacterial osteomyelitis in humans. Current opinion in rheumatology. 2018;30(5):521–5.

14. Chitu V, Ferguson PJ, de Bruijn R, Schlueter AJ, Ochoa LA, Waldschmidt TJ, et al. Primed innate immunity leads to autoinflammatory disease in PSTPIP2-deficient cmo mice. Blood. 2009;114(12):2497–505.

15. Liu L, Wen Q, Gong R, Gilles L, Stankiewicz MJ, Li W, et al. PSTPIP2 dysregulation contributes to aberrant terminal differentiation in GATA-1-deficient megakaryocytes by activating LYN. Cell death & disease. 2014;5:e988.

16. Cassel SL, Janczy JR, Bing X, Wilson SP, Olivier AK, Otero JE, et al. Inflammasome-independent IL-1beta mediates autoinflammatory disease in Pstpip2-deficient mice. Proceedings of the National Academy of Sciences of the United States of America. 2014;111(3):1072–7.

17. Lukens JR, Gross JM, Calabrese C, Iwakura Y, Lamkanfi M, Vogel P, et al. Critical role for inflammasome independent IL-1β production in osteomyelitis. Proceedings of the National Academy of Sciences of the United States of America. 2014;111(3):1066–71.

18. Chitu V, Pixley FJ, Macaluso F, Larson DR, Condeelis J, Yeung YG, et al. The PCH family member MAYP/PSTPIP2 directly regulates F-actin bundling and enhances filopodia formation and motility in macrophages. Mol Biol Cell. 2005;16(6):2947–59.

19. Tsujita K, Kondo A, Kurisu S, Hasegawa J, Itoh T, Takenawa T. Antagonistic regulation of F-BAR protein assemblies controls actin polymerization during podosome formation. J Cell Sci. 2013;126(Pt 10):2267–78.

20. Netea MG, van de Veerdonk FL, van der Meer JW, Dinarello CA, Joosten LA. Inflammasome-independent regulation of IL-1-family cytokines. Annual review of immunology. 2015;33:49–77.

21. Lukens JR, Gurung P, Vogel P, Johnson GR, Carter RA, McGoldrick DJ, et al. Dietary modulation of the microbiome affects autoinflammatory disease. Nature. 2014;516(7530):246–9.

22. Chitu V, Nacu V, Charles JF, Henne WM, McMahon HT, Nandi S, et al. PSTPIP2 deficiency in mice causes osteopenia and increased differentiation of multipotent myeloid precursors into osteoclasts. Blood. 2012;120(15):3126–35.

23. Itoh T, Erdmann KS, Roux A, Habermann B, Werner H, De Camilli P. Dynamin and the actin cytoskeleton cooperatively regulate plasma membrane invagination by BAR and F-BAR proteins. Dev Cell. 2005;9(6):791–804.

24. Tsujita K, Suetsugu S, Sasaki N, Furutani M, Oikawa T, Takenawa T. Coordination between the actin cytoskeleton and membrane deformation by a novel membrane tubulation domain of PCH proteins is involved in endocytosis. J Cell Biol. 2006;172(2):269–79.

25. Wu Y, Dowbenko D, Lasky LA. PSTPIP 2, a second tyrosine phosphorylated, cytoskeletal-associated protein that binds a PEST-type protein-tyrosine phosphatases. The Journal of biological chemistry. 1988;273(46):30487–96.

26. Drobek A, Kralova J, Skopcova T, Kucova M, Novak P, Angelisova P, et al. PSTPIP2, a Protein Associated with Autoinflammatory Disease, Interacts with Inhibitory Enzymes SHIP1 and Csk. J Immunol. 2015;195(7):3416–26.

27. Yeung YG, Soldera S, Stanley ER. A novel macrophage actin-associated protein (MAYP) is tyrosine-phosphorylated following colony stimulating factor-1 stimulation. J Biol Chem. 1998;273(46):30638–42.

28. Cloutier JF, Veillette A. Association of inhibitory tyrosine protein kinase p50csk with protein tyrosine phosphatase PEP in T cells and other hemopoietic cells. EMBO J. 1996;15(18):4909–18.

29. Cloutier JF, Veillette A. Cooperative inhibition of T-cell antigen receptor signaling by a complex between a kinase and a phosphatase. J Exp Med. 1999;189(1):111–21.

30. Rauh MJ, Sly LM, Kalesnikoff J, Hughes MR, Cao LP, Lam V, et al. The role of SHIP1 in macrophage programming and activation. Biochemical Society transactions. 2004;32(Pt 5):785–8.

31. Huber M, Gibbs BF. SHIP1 and the negative control of mast cell/basophil activation by supra-optimal antigen concentrations. Molecular immunology. 2015;63(1):32–7.

32. Tsai M, Grimbaldeston M, Galli SJ. Mast cells and immunoregulation/immunomodulation. Adv Exp Med Biol. 2011;716:186–211.

33. Wernersson S, Pejler G. Mast cell secretory granules: armed for battle. Nat Rev Immunol. 2014;14(7):478–94.

34. De Filippo K, Dudeck A, Hasenberg M, Nye E, van Rooijen N, Hartmann K, et al. Mast cell and macrophage chemokines CXCL1/CXCL2 control the early stage of neutrophil recruitment during tissue inflammation. Blood. 2013;121(24):4930–7.

35. Haeggstrom JZ, Funk CD. Lipoxygenase and leukotriene pathways: biochemistry, biology, and roles in disease. Chemical reviews. 2011;111(10):5866–98.

36. Satpathy SR, Jala VR, Bodduluri SR, Krishnan E, Hegde B, Hoyle GW, et al. Crystalline silica-induced leukotriene B4-dependent inflammation promotes lung tumour growth. Nature communications. 2015;6:7064.

37. Dudeck A, Dudeck J, Scholten J, Petzold A, Surianarayanan S, Kohler A, et al. Mast cells are key promoters of contact allergy that mediate the adjuvant effects of haptens. Immunity. 2011;34(6):973–84.

38. Dudeck J, Ghouse SM, Lehmann CH, Hoppe A, Schubert N, Nedospasov SA, et al. MastCell-Derived TNF Amplifies CD8(+) Dendritic Cell Functionality and CD8(+) T Cell Priming. Cell reports. 2015;13(2):399–411.

39. Schubert N, Dudeck J, Liu P, Karutz A, Speier S, Maurer M, et al. Mast cell promotion of T cell-driven antigen-induced arthritis despite being dispensable for antibody-induced arthritis in which T cells are bypassed. Arthritis & rheumatology. 2015;67(4):903–13.

40. Feldkamp LA, Davis LC, Kress JW. Practical Cone-Beam Algorithm. J Opt Soc Am A. 1984;1(6):612–9.

41. Doube M, Klosowski MM, Arganda-Carreras I, Cordelieres FP, Dougherty RP, Jackson JS, et al. BoneJ: Free and extensible bone image analysis in ImageJ. Bone. 2010;47(6):1076–9.

42. Sharma N, Kumar V, Everingham S, Mali RS, Kapur R, Zeng LF, et al. SH2 domain-containing phosphatase 2 is a critical regulator of connective tissue mast cell survival and homeostasis in mice. Mol Cell Biol. 2012;32(14):2653–63.

43. Lukens JR, Gross JM, Calabrese C, Iwakura Y, Lamkanfi M, Vogel P, et al. Critical role for inflammasome-independent IL-1beta production in osteomyelitis. Proc Natl Acad Sci U S A. 2014;111(3):1066–71.

44. Kroner J, Kovtun A, Kemmler J, Messmann JJ, Strauss G, Seitz S, et al. Mast Cells Are Critical Regulators of Bone Fracture-Induced Inflammation and Osteoclast Formation and Activity. Journal of bone and mineral research: the official journal of the American Society for Bone and Mineral Research. 2017.

45. Caughey GH. Mast cell tryptases and chymases in inflammation and host defense. Immunological Reviews. 2007;217:141–54.

46. Dong X, Chen J, Zhang Y, Cen Y. Mast cell chymase promotes cell proliferation and expression of certain cytokines in a dose-dependent manner. Molecular medicine reports. 2012;5(6):1487–90.

47. Zhang H, Huang Y, Wang S, Fu R, Guo C, Wang H, et al. Myeloid-derived suppressor cells contribute to bone erosion in collagen-induced arthritis by differentiating to osteoclasts. Journal of autoimmunity. 2015;65:82–9.

48. Sawant A, Ponnazhagan S. Myeloid-derived suppressor cells as osteoclast progenitors: a novel target for controlling osteolytic bone metastasis. Cancer Res. 2013;73(15):4606–10.

49. Fujii W, Ashihara E, Hirai H, Nagahara H, Kajitani N, Fujioka K, et al. Myeloid-derived suppressor cells play crucial roles in the regulation of mouse collagen-induced arthritis. J Immunol. 2013;191(3):1073–81.

50. Zhang H, Wang S, Huang Y, Wang H, Zhao J, Gaskin F, et al. Myeloid-derived suppressor cells are proinflammatory and regulate collagen-induced arthritis through manipulating Th17 cell differentiation. Clinical immunology. 2015;157(2):175–86.

51. Hegde VL, Singh UP, Nagarkatti PS, Nagarkatti M. Critical Role of Mast Cells and Peroxisome Proliferator-Activated Receptor gamma in the Induction of Myeloid-Derived Suppressor Cells by Marijuana Cannabidiol In Vivo. J Immunol. 2015; 194(11):5211–22.

52. Feyerabend TB, Weiser A, Tietz A, Stassen M, Harris N, Kopf M, et al. Cre-mediated cell ablation contests mast cell contribution in models of antibody-and T cell-mediated autoimmunity. Immunity. 2011;35(5):832–44.

53. Dawicki W, Jawdat DW, Xu N, Marshall JS. Mast cells, histamine, and IL-6 regulate the selective influx of dendritic cell subsets into an inflamed lymph node. J Immunol. 2010;184(4):2116–23.

54. Oldford SA, Haidl ID, Howatt MA, Leiva CA, Johnston B, Marshall JS. A critical role for mast cells and mast cell-derived IL-6 in TLR2-mediated inhibition of tumor growth. J Immunol. 2010;185(11):7067–76.

55. Sugitharini V, Shahana P, Prema A, Berla Thangam E. TLR2 and TLR4 co-activation utilizes distinct signaling pathways for the production of Th1/Th2/Th17 cytokines in neonatal immune cells. Cytokine. 2016;85:191–200.

56. Mizutani H, Schechter N, Lazarus G, Black RA, Kupper TS. Rapid and specific conversion of precursor interleukin 1 beta (IL-1 beta) to an active IL-1 species by human mast cell chymase. J Exp Med. 1991;174(4):821–5.

57. Bonnekoh H, Scheffel J, Kambe N, Krause K. The role of mast cells in autoinflammation. Immunological reviews. 2018;282(1):265–75.

